# Dissection of the *Fgf8* regulatory landscape by *in vivo* CRISPR-editing reveals extensive inter- and intra-enhancer redundancy

**DOI:** 10.1101/2020.03.03.966796

**Authors:** A. Hörnblad, K. Langenfeld, S. Bastide, F. Langa Vives, F. Spitz

## Abstract

Developmental genes are often regulated by multiple elements with overlapping activity. Yet, in most cases, the relative function of those elements and their contribution to endogenous gene expression remain uncharacterized. Illustrating this situation, distinct sets of enhancers have been proposed to direct *Fgf8* in the limb apical ectodermal ridge (AER) and the midbrain-hindbrain boundary (MHB). Using *in vivo* CRISPR/Cas9 genome engineering, we functionally dissect this complex regulatory ensemble and demonstrate two distinct regulatory logics. In the AER, the control of *Fgf8* expression appears extremely distributed between different enhancers. In contrast, in the MHB, one of the three active enhancers is essential while the other two are dispensable. Further dissection of the essential MHB enhancer revealed another layer of redundancy and identified two sub-parts required independently for *Fgf8* expression and formation of midbrain and cerebellar structures. Interestingly, cross-species transgenic analysis of this enhancer suggests changes of the organisation of this essential regulatory node in the vertebrate lineage.

## Introduction

A fundamental feature of animal development is the dynamic and highly reproducible spatiotemporal expression of many key genes. This spatial and temporal specificity is coordinated through the actions of *cis*-regulatory elements that can reside very far (up to Mb) from their target genes and even be located within neighbouring genes ^1–6^. Transgenic studies have been important to identify enhancer sequences with regulatory activity in the genome ^7^, but with a low throughput. More recently, next generation sequencing approaches such as chromosome conformation capture, ChIP-seq, DNAse-seq and ATAC-seq allowed for more comprehensive identification of candidate regulatory regions ^1,2,4,8^. These studies have demonstrated that the regulatory architecture of developmental genes is complex: it frequently includes multiple regulatory elements, dispersed over large genomic regions ^9^, that often display overlapping and/or redundant activity. As useful they are, a strong limitation of these approaches is that they do not determine how important those elements are for gene expression. Indeed, it happens frequently that enhancers with strong transgenic activities have a surprisingly minor function *in vivo* in the control of their endogenous gene ^10–13^. Because of this difference between function and activity, there is an urgent need to develop strategies to characterize the biological function of non-coding regulatory elements *in vivo* and *in situ*. Traditional gene targeting approaches have demonstrated the functional importance of individual enhancers, but the throughput of these techniques is relatively low ^14–16^. Here, we deployed a Crispr/Cas9 *in vivo* genome-engineering approach to systematically dissect the functional importance of individual enhancers as well as their intrinsic logic *in vivo*, using the *Fgf8* locus as a model system.

FGF8 is a secreted signalling molecule with a highly dynamic gene expression pattern during development. It is essential for the normal development of the brain, craniofacial skeleton, limbs, and various other organs ^17–22^. FGF8 is the key molecule for the formation and activity of the isthmic organizer (IsO) located at the border between the mesencephalon and metencephalon ^23–25^ and that plays essential roles for patterning the midbrain and cerebellum ^17,26^. Targeted deletion of *Fgf8* in the MHB leads to down-regulation of MHB markers and subsequent loss of the midbrain and anterior hindbrain ^26^. In the limb, *Fgf8* is expressed in apical ectodermal ridge (AER), at the distal tip of the limb bud. Absence of *Fgf8* leads to aberrant proximo-distal and anterior-posterior patterning, increased apoptosis in the limb bud and subsequent loss or hypoplasia of specific skeletal ^20,21^.

Although the consequences of *Fgf8* down-regulation in the MHB and AER have been well characterized ^20,21,26–28^, less is known about the regulatory elements directing *Fgf8* expression in these structures. In a recent study, we characterized a 200kb region forming the *Fgf8* regulatory landscape and identified three enhancers with the potential to drive expression in the mouse MHB and five enhancers that could drive expression in the limb AER ^6^. The MHB enhancers are highly conserved from fish to mammals and two of them have indeed been identified as potential drivers of *Fgf8* expression also in the zebrafish MHB ^6,29–31^. The limb enhancers show a more diverse degree of conservation but all of them are conserved at least from amniotes to mammals ^6^.

In this study we address the *in vivo* contribution of these two sets of enhancers to *Fgf8* expression in the limb and the MHB, respectively. Using CRISPR/Cas9 genome editing we demonstrate extensive redundancy between enhancers in the limb, while in the MHB, one distant primary enhancer is essential for *Fgf8* expression. We further dissect the main MHB enhancer extensively to identify its functional units and define two essential subunits required for its function. Intriguingly, although deletion of only 37bp is enough to abrogate the regulatory potential of this enhancer and cause loss of midbrain and cerebellar structures, we also reveal widespread functional redundancy within this essential enhancer. Furthermore, we demonstrate that albeit sequence conservation predicts similar enhancer activity in fish and mouse, the functional subunits of the enhancer appear to have diverged and reorganized their regulatory logic.

## Results

### Extensive regulatory redundancy for *Fgf8* expression in the limb

A previous study identified a set of putative limb and MHB enhancers in the *Fgf8* locus with the potential to drive gene expression in these tissues ^6^ (Fig1A). In order to investigate their *in vivo* role, we generated mice with targeted deletions of each individual enhancer as well as compound deletions of the two proximal MHB enhancers. To this end we performed zygote injections of Cas9 mRNA and two chimeric gRNAs flanking the regions of interest (FigS1, TableS1 and methods, *in vivo* deletion efficiency ranging from 4 to 40% in born pups). We assessed the consequence of these enhancer deletions in hemizygous condition over *Fgf8* null alleles (either Fgf8^null/+ 17^ or DEL(P-F8) ^6^).

**Fig1.**
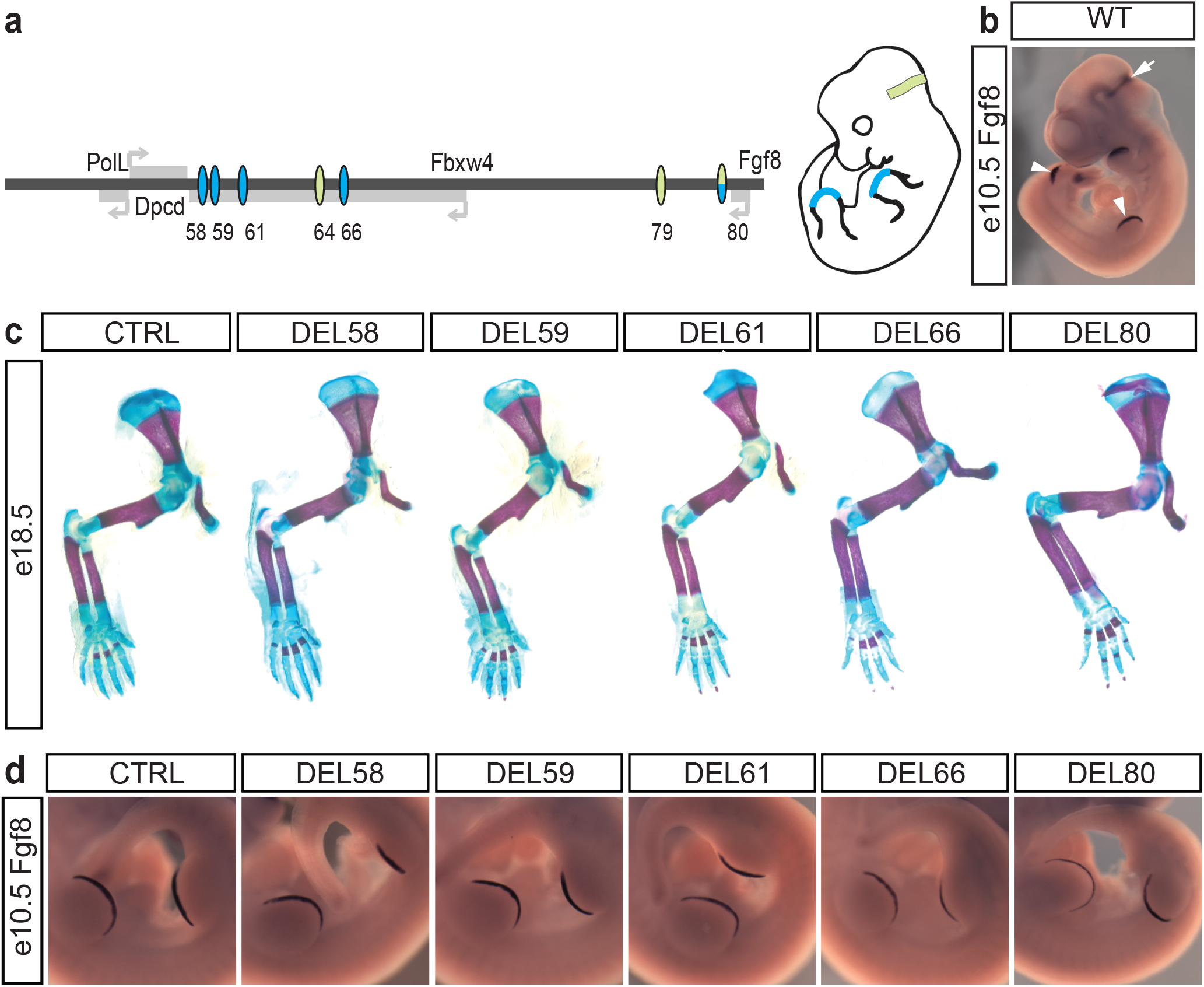
The limb AER elicits extensive regulatory redundancy. (**A**) Schematic representation of the two sets of conserved elements directing expression in the AER (blue), and in the MHB (green). (**B**) In situ hybridization with riboprobe against *Fgf8* mRNA. Arrowheads and arrow indicate AER and MHB, respectively. (**C**) Photomicrograph of alizarin-red/alcian-blue stained e18.5 forelimbs from control and AER enhancer deletion embryos. (**D**) In situ hybridization of control and AER enhancer deletion embryos at e10.5 with riboprobe against *Fgf8*. All mutant embryos display expression patterns indistinguishable from their littermate controls.

For the limb, four enhancers (CE58, CE59, CE61, CE66) are spread within a 40kb region in the introns of the neighbouring *Fbxw4* gene while only CE80 is located in the proximity of *Fgf8* (Fig1A). Previous experiments had demonstrated that mice carrying a deletion of the region containing the four distal enhancers abolishes limb *Fgf8* expression and causes similar defects to the conditional ablation of *Fgf8* in the limb. In contrast, all the mutants that we generated carrying single deletions of these putative enhancers were healthy and had limbs indistinguishable from their control littermates. These results were confirmed in more detail by skeletal preparations of e18.5 embryos (Fig1C). We also analysed the expression pattern of *Fgf8* at e10.5 using *in situ* hybridisation. At this stage *Fgf8* is strongly expressed in the morphologically well-defined AER of both the forelimb and the hindlimb (Fig1B). The AER expression pattern displayed by embryos carrying enhancer deletions was indistinguishable from their control littermates (Fig1D).

To further confirm this, we performed quantitative RT-qPCR analysis on dissected e10.5 forelimbs of three deletion lines (DEL58, DEL61, DEL80, corresponding to deeply evolutionary conserved enhancers) and failed to detect significant change in *Fgf8* gene expression levels or in other limb patterning genes, which could have indicated compensatory effects (FigS2). Thus, from a pure functional viewpoint, each of those enhancers appears dispensable for the expression of *Fgf8* and subsequent development of the limb. Taken together, this demonstrates that the regulatory system that controls *Fgf8* limb expression *in vivo* is highly modular and displays extensive regulatory redundancy.

### A distant *Fgf8* enhancer is required for formation of the midbrain and cerebellum

In the MHB, two of the putative enhancers (CE79 and CE80) are located within a 20kb region downstream of *Fgf8*, while the third one (CE64) is located at a distance of 120kb within an intron of the neighbouring gene *Fbxw4* (Fig1A). Using CRISPR/Cas9 zygote injections, we generated mice carrying single deletions of these enhancers as well as the double deletion of CE79 and CE80. We found no morphological differences between the DEL79, DEL80 or the compound DEL79-80 animals and their control littermates that could be detected macroscopically in the brain. In contrast, DEL64 mice display a complete absence of midbrain and cerebellar structures visible at e18.5, phenocopying the conditional KO of Fgf8 in the MHB ^26^. A more detailed analysis of e18.5 brains using optical projection tomography (OPT) demonstrates the complete loss of superior colliculus, inferior colliculus, isthmus and cerebellum in the DEL64 mutants (Fig2, VideoS1). These analyses also confirmed the normal appearance of these structures in the DEL79, DEL80, and DEL79-80 mutants (Fig2, VideoS2-4). In summary, of the three MHB enhancers, only CE64 is essential for proper development of the MHB. Despite the sensitivity of the MHB-derived structures to mild-reduction of Fgf8-signalling from the IsO, which could result in various degrees of hypoplasia ^17,28^, the two proximal enhancers 79 and 80 appear dispensable for the development of those structures.

**Fig2.**
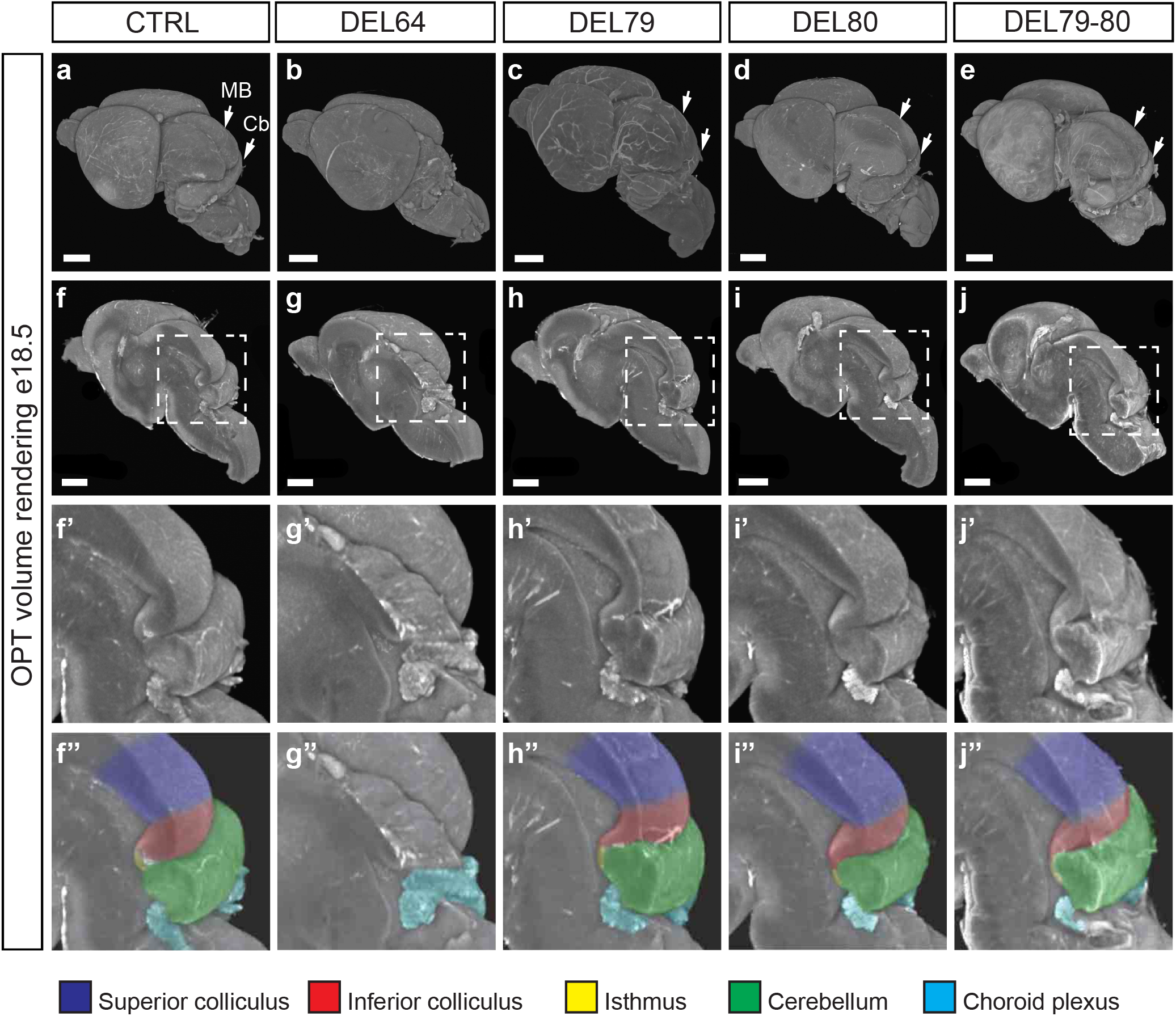
One main enhancer is required for Fgf8 expression in the MHB. (**A** to **J’’**) OPT generated volume renderings of e18.5 brains from control (**A**, **F**, **F’**, **F’’**), DEL64 (**B**, **G**, **G’**, **G’’**), DEL79 (**C**, **H**, **H’**, **H’’**), DEL80 (**D**, **I**, **I’**, **I’’**) and DEL79-80 (**E**, **J**, **J’**, **J’’**) mutants. Signal is based on tissue autoflourescence. Control brains, DEL79, DEL80 and DEL79-80 mutants display a well-developed midbrain and cerebellar anlage while DEL64 brains display severe hypoplasia of midbrain and cerebellum. (**F** to **J**) Midsagittal digital dissection reveal complete loss of all MHB derived structures in the DEL64 mutant. (**F’** to **J’’**) Close-up of boxed area in (**F** to **J**). Brains have been pseudocolored in (**F’’** to **J’’**): dark blue – superior colliculus; red – inferior colliculus; yellow – isthmus; green – cerebellum; light blue – choroid plexus. Scalebar in (**A-J**) is 1mm. MB, midbrain; Cb, cerebellum.

### Deletion of CE64 completely abolishes Fgf8 expression in the MHB

We further explored the spatial expression of Fgf8 at e10.5 in all the generated MHB mutants (Fig3A-F). At this time point in development, the expression of *Fgf8* has been narrowed down to a sharply delimited band of cells at the border between the midbrain and anterior hindbrain. In the DEL64 embryos *Fgf8* expression was completely absent in the MHB and the morphology of these embryos already revealed the absence of a large portion of the midbrain (Fig3B). In DEL79, DEL80 and DEL79-80 embryos, *Fgf8* expression pattern and signal strength were similar to control embryos. Next, we performed *in situ* hybridisation analysis of *Fgf8* expression at the earliest stage of expression, e8.25, in DEL64 embryos. These analyses revealed a complete lack of *Fgf8* expression also in the initial expression phase (Fig3G-H). This indicates that CE64 is required and sufficient for proper initiation of *Fgf8* expression as well as subsequent maintenance.

**Fig3.**
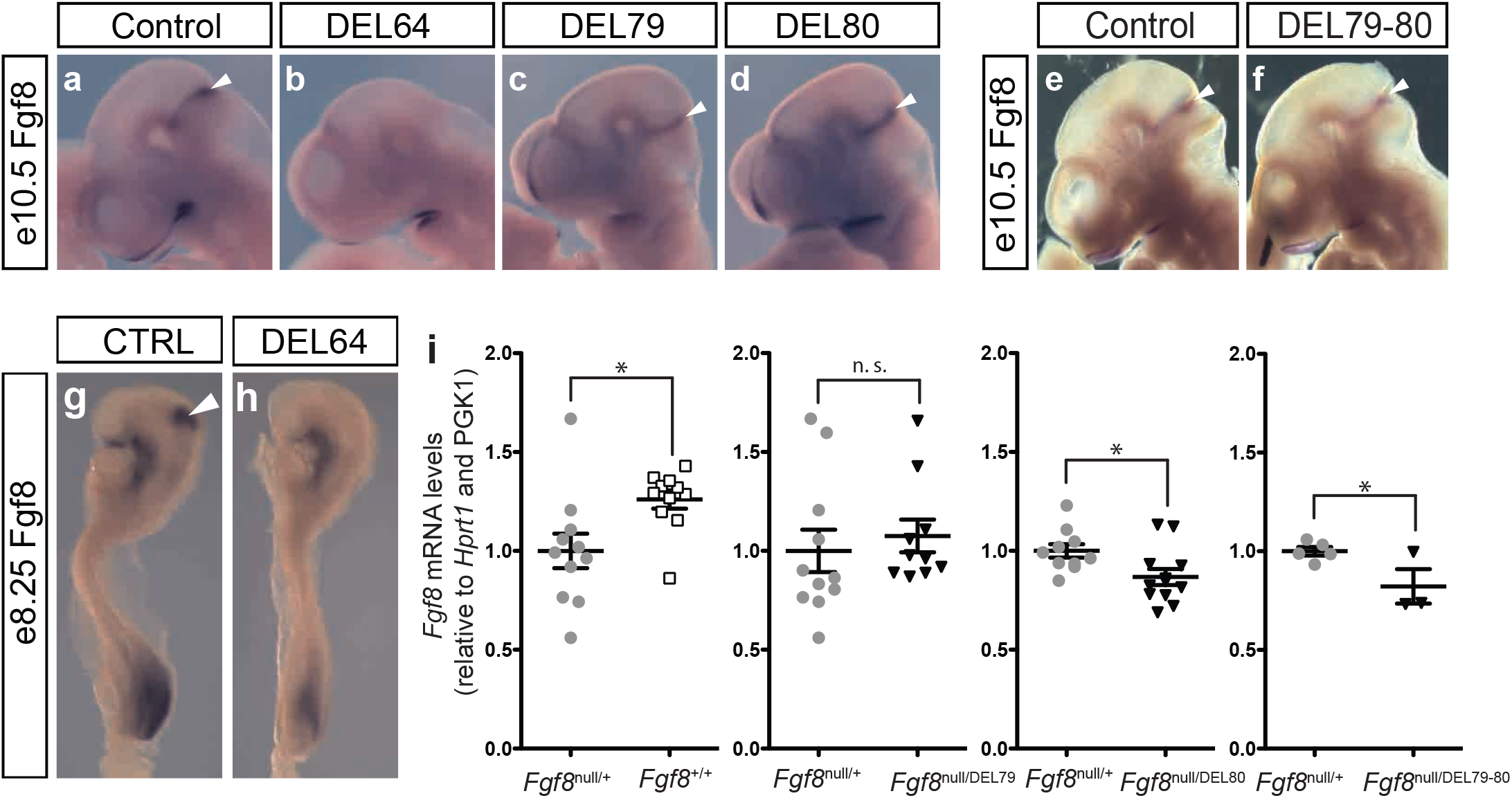
CE64 is required and sufficient for initiation and maintenance of *Fgf8* expression in the MHB. (**A** to **F**) *In situ* hybridization against *Fgf8* mRNA in e10.5 embryos from control (**A, E**), DEL64 (**B**), DEL79 (**C**), DEL80 (**D**) and DEL79-80 (**F**) mutants. Note the complete lack of Fgf8 expression in the MHB of DEL64 (**B**) embryos as compared to controls (**A**). Brain tissue is markedly reduced already at this stage in DEL64 embryos (**B**). (**G** and **H**) In situ hybridization of control and DEL64 enhancer deletion embryos at e8.25 with riboprobe against *Fgf8*. No expression is detected in the DEL64 embryos. (**I**) Relative expression of *Fgf8* mRNA levels in dissected MHB region from WT (n=11), DEL79 (n=10), DEL80 (n=12), and DEL79-80 (n=3) e10.5 embryos as compared to Fgf8^null/+^ (n=11, 11, 10, 5) control littermates. Individual data point, mean±SEM are indicated. *p<0.05, n. s. = not significant (two-tailed Student’s *t*-test).

Although *in situ* hybridisation revealed similar expression patterns between DEL79, DEL80 and DEL79-80 mutants as compared to control embryos, we sought to assess potential subtle quantitative changes in the expression levels. For this, we performed RT-qPCR on dissected MHB region from e10.5 embryos, for *Fgf8* and a set of genes known to be involved in this gene regulatory network.

Firstly, we noticed that mice heterozygous for a null *Fgf8* allele only showed a mild reduction of *Fgf8*. Instead of an expected 50% reduction, we measured that *Fgf8* expression in *Fgf8*^*null*/*+*^ was 79% of wild-type level in the MHB, and 68% in the limb (Fig3I, FigS2). This limited impact suggests that *Fgf8* expression is maintained, at least partially, by feedback mechanisms. In *Fgf8*^*null*/*+*^ MHB, we found a decreased expression of *Spry2* and *Dusp6* (FigS3), two downstream targets of *Fgf8* that are part of negative feedback loops for Fgf-signalling ^32,33^, suggesting that this circuit could account for sustained expression upon *Fgf8* gene dosage reduction.

Taking this potential compensation into account, in the enhancer deletion alleles we could detect a mild but significant decrease in expression of *Fgf8* as compared to the control animals for the DEL80 as well as the compound DEL79-80 (Fig3I). This decrease was accompanied by a small but significant decrease in *Dusp6, En1, En2, Fgf17, Spry1* for DEL80, and *Dusp6, Etv4, Fgfr1, Lmx1b, Pax2, Sp8, Spry1, and Spry2* for DEL79-80 (FigS3). The general tendency in these mutants is a minor down-regulation of the genes in the MHB regulatory network. Notably, the expression profiles of DEL80 and DEL79-80 tend to overlap and may indicate that most of the effects seen in the compound mutant is due to the deletion of CE80. In contrast, most genes investigated in the DEL79 embryos display a tendency to minor up-regulation that is significant for *Fgf17, Lmx1b, Otx2, Pax2, Pax6, Spry1*, and *Spry2* (FigS3). Taken together, these data demonstrate only minor contribution of CE79 and CE80 to *Fgf8* gene expression and hence underline the essential role of the main CE64 enhancer in MHB development. Thus, CE64 appears as the main enhancer of *Fgf8* expression in the MHB, that it is required and sufficient for the initiation of *Fgf8* expression, while both CE79 and CE80 are dispensable for MHB patterning.

### *In vivo* CRISPR/*Cas9* screen identifies two distinct subunits required for CE64 enhancer function

Given the crucial role of CE64 for the expression of *Fgf8* in the MHB we aimed to dissect how the regulatory logic of this enhancer is composed *in vivo*. To this end, we injected a new set of CRISPR gRNAs in different combinations together with *Cas9* mRNA in oocytes that had been *in vitro* fertilized using sperm from males heterozygous for the DEL(P-F8) allele (Fig4A). The fact that 50% of injected embryos carried DEL(P-F8) increased the yield of “informative” embryos (only deletion of one enhancer copy is required) and facilitated their unambiguous identification (reduced possible mosaicism). As disruption of CE64 function leads to a severe hypoplasia of the midbrain and cerebellum, we could directly screen F0 embryos at e18.5 for lack of these tissues (Fig4B) and identify regions essential to CE64 function in embryos carrying DEL(P-F8) compound or homozygous targeted deletions. Using this system, we produced and analysed a large collection of deletions spanning different regions of CE64, performing *in vivo*, at the endogenous locus, the type of “enhancer-bashing” experiments that are typically carried out on out-of-context transgenic assays. All embryos produced were genotyped by PCR for targeted deletions and the breakpoints were sequenced. In addition, the embryos carrying deletions were genotyped with primers internal to the identified deletions in order to discard embryos carrying WT alleles due to mosaicism (Fig4C). In all, we identified 39 informative alleles (TableS2).

**Fig4.**
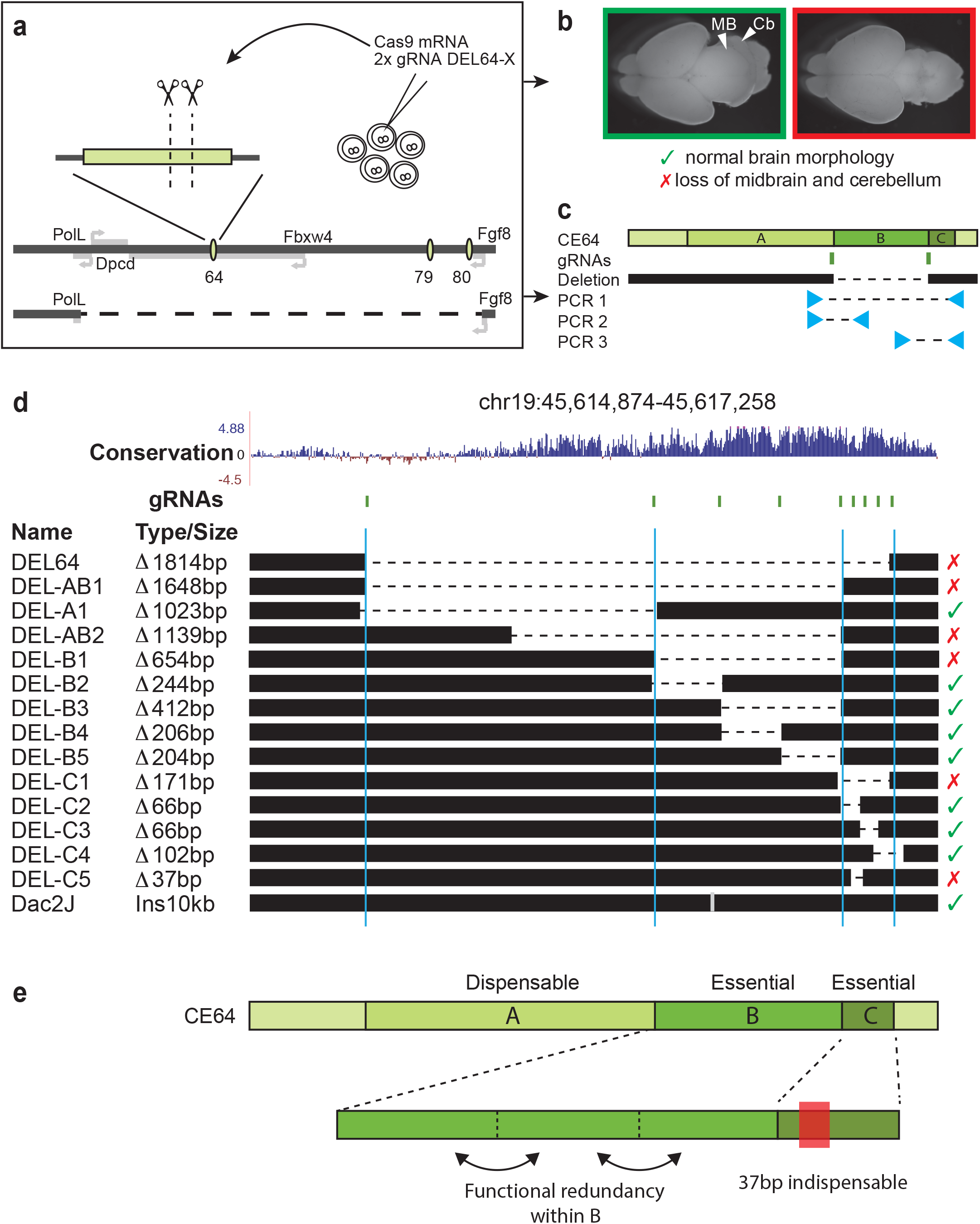
Two distinct subunits with internal redundancy are required for CE64 function. (**A**) Schematic representation of the CRISPR screen setup. Oocytes were fertilized with sperm carrying one DEL(P-F8) allele (bottom) and 2 gRNAs were simultaneously injected with *Cas9* mRNA. (**B**) Brain morphology of F0 embryos was examined at e18.5 and all embryos were genotyped according to strategy in (**C**) to identify breakpoints and possible mosaicism. (**D**) Representation of the panel of deletions generated. The *in vivo* CRISPR/*Cas9* screen defined two indispensable subunits of CE64 (DEL-B and DEL-C). Removing overlapping bits within these units (DEL-B2 through DEL-B5, DEL-C2 through DEL-C4) does not provoke any phenotype. The smallest deletion causing lack of MHB derived structures is merely 37bp (DEL-C5). Red cross indicates loss of MHB derived tissues and green tick indicates normal brain morphology. (**E**) Schematic representation of the functional units of CE64. Both 64-B and 64-C are indispensable for CE64 function, while 64-A is not required. Functional redundancy is encoded within these subunits although a deletion of only 37bp is enough to abrogate the function of 64-C.

This extensive panel of deletions allowed us to define three distinct elements in CE64, of which one is dispensable (64-A in Fig4E) and two (64-B and 64-C in Fig4E) are essential and required for proper enhancer function. Deleting any of the two essential regions 64-B or 64-C is sufficient to completely abrogate the development of the midbrain and anterior hindbrain region. Of these essential subunits, 64-B spans a region of approximately 700bp that is highly conserved among vertebrates (Fig4D, Fig6A). Intriguingly, deletions of sub-regions in 64-B demonstrated that considerable functional redundancy exists within this subunit. In fact, deleting two-thirds of 64-B is not sufficient to abolish proper midbrain and cerebellum formation (DEL-B3 in Fig4D) and any one third of this subunit is dispensable for its function (DEL-B2, DEL-B4, DEL-B5 in Fig4D). Of note, a 10kb insertion of a MusD retroelement as observed in the Dac2J strain (Fig4D), appear to have no impact on MHB development (Tugce Aktas, in preparation). Therefore, it seems that the regulatory information embedded in 64-B is modular and spread across the element, rather than organised as one continuous regulatory unit. Subunit 64-C is only 180bp long and located on the most telomeric side of CE64. It is conserved in tetrapods but not in fish. Consecutive deletions of sub-regions in 64-C do not cause any phenotype (DEL-C2, DEL-C3, DEL-C4 in Fig4, Fig6D), but remarkably, the deletion of merely 37bp in 64-C at the junction between 64-C2 and 64-C3 is sufficient to completely abrogate CE64 function (DEL-C5 in Fig4A, Fig 6D). This indicates that the 37bp contains at least two critical, yet redundant elements.

### The functional subunits of CE64 are interdependent

Next, we asked if 64-B and 64-C differ in their regulatory potential by performing transient transgenesis of a reporter construct carrying either CE64, 64-B or CE64 lacking 64-B or 64-C, respectively. The sequences were cloned upstream of a minimal promoter that by itself does not drive expression of the LacZ reporter gene. As expected, Tg(CE64) recapitulated the expression pattern published for CE64 in 3 out of 4 transgenic embryos (Fig5C and FigS4, for comparison see Marinic *et al.*) ^6^. However, for both Tg(64-DEL-B) and the Tg(64-DEL-C), no expression was detected in the MHB (0/8 and 0/3 embryos respectively) (Fig5C, FigS4). Interestingly, some of the Tg(DEL-B) embryos (4/8) displayed a reproducible reporter expression in the anterior hindbrain (FigS4). This may indicate that 64-C has an intrinsic regulatory potential that is independent of 64-B for expression *per se* but which spatial position is shifted in presence of 64-B. In contrast, 64-B does not appear to have any autonomous activity in e10.5 embryos (0/4 embryos) (Fig5C, FigS4). Taken together, the transgenic assays indicate that although both 64-B and 64-C are required for the function of CE64, their intrinsic properties are not sufficient to drive spatial expression in the MHB on their own.

**Fig5.**
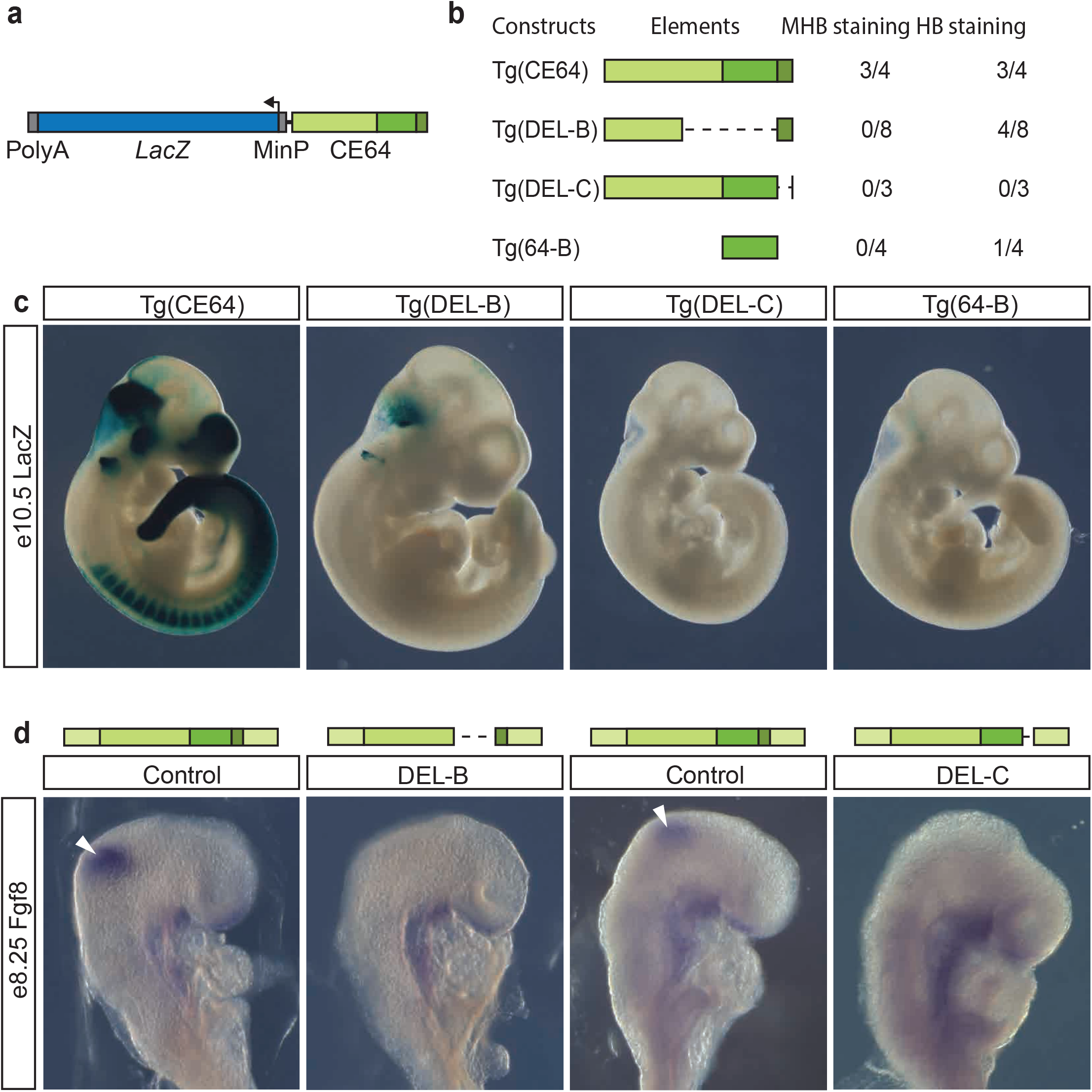
The function of 64-B and 64-C is interdependent. (**A**) Schematic of transgenic reporter construct containing *LacZ*, a minimal promoter and the putative enhancer sequence of interest. (**B**) Table of constructs used for transgenesis, the CE64 subunits included and the number of embryos displaying reporter expression for each construct. (**C**) Photomicrographs of representative embryos stained for *LacZ* activity. Reporter expression in the MHB is only detected in wt Tg(CE64) embryos. Note the staining that is present in the anterior hindbrain of some of the Tg(DEL-B) embryos. Tg(DEL-C) and Tg(64-B) do not manifest any reproducible reporter expression. (**D**) *In situ* hybridization of control, DEL-B, and DEL-C deletion embryos at e8.25 with riboprobe against *Fgf8*. Expression is undetectable in both enhancer subunit deletions. Blue box indicates *LacZ* reporter gene in (**A**). In all panels, light green, green and dark green boxes indicate 64-A, 64-B, and 64-C subunits, respectively.

The transient transgenesis and CRISPR/*Cas9* deletion screen of F0 progeny at late stages of embryonic development precluded direct analysis of *Fgf8* gene expression at the time when the MHB patterns the prospective midbrain and hindbrain. In order to assess the contribution of each of the CE64 sub-units to *Fgf8* expression in the MHB during early stages we generated mouse lines carrying deletions of 64-B and 64-C. As expected, both lines completely lack midbrain and cerebellum at e18.5, confirming the results from the embryonic screen. Expression analysis by in situ hybridisation at e8.25 demonstrated that in both mutants, *Fgf8* gene expression fails to initiate and is completely absent from the MHB region (Fig5D). Altogether, these data demonstrate that the functional elements of CE64 are units that have reciprocal dependency in order to mediate proper regulatory input to *Fgf8*.

### Evolutionary conservation of CE64 sequence versus functional organisation

Conservation is a good predictor for identifying regulatory regions in the genome and a previous study has shown that the zebrafish region orthologous to CE64 can drive expression in the zebrafish MHB (dr10 in ref. ^31^). Intriguingly, our functional analysis in mouse of CE64 sub-regions identified an essential part of the enhancer (64-C) that is not conserved in fish (Fig6A). In addition, transgenic analysis showed that the conserved 64-B element is unable to drive expression in the MHB by its own (Fig5C). We therefore asked whether the orthologous region in fish could drive MHB expression in the mouse. To this end, we cloned CE64 from spotted gar, a species that is closer to mouse and humans in the vertebrate lineage and has not undergone the genome duplication that the teleost lineage has. Remarkably, the 350bp sequence from spotted gar could drive expression in the MHB region in 4 out of 4 embryos (Fig6B, FigS4), despite lacking a region orthologous to the mouse 64-C. The expression did not completely reproduce the expression of the full CE64 but was restricted to the MHB and dorsal part of the anterior hindbrain. It is also noteworthy that the zebrafish dr10 enhancer recapitulates the broad activity of mouse CE64 in the MHB region (as well as in the forebrain and tail bud), in the zebrafish transcriptional context. This raises the question to whether non-conserved sequences outside the 350bp core enhancer may encode additional information that would further increase the similarity in regulatory potential to mouse CE64.

**Fig6.**
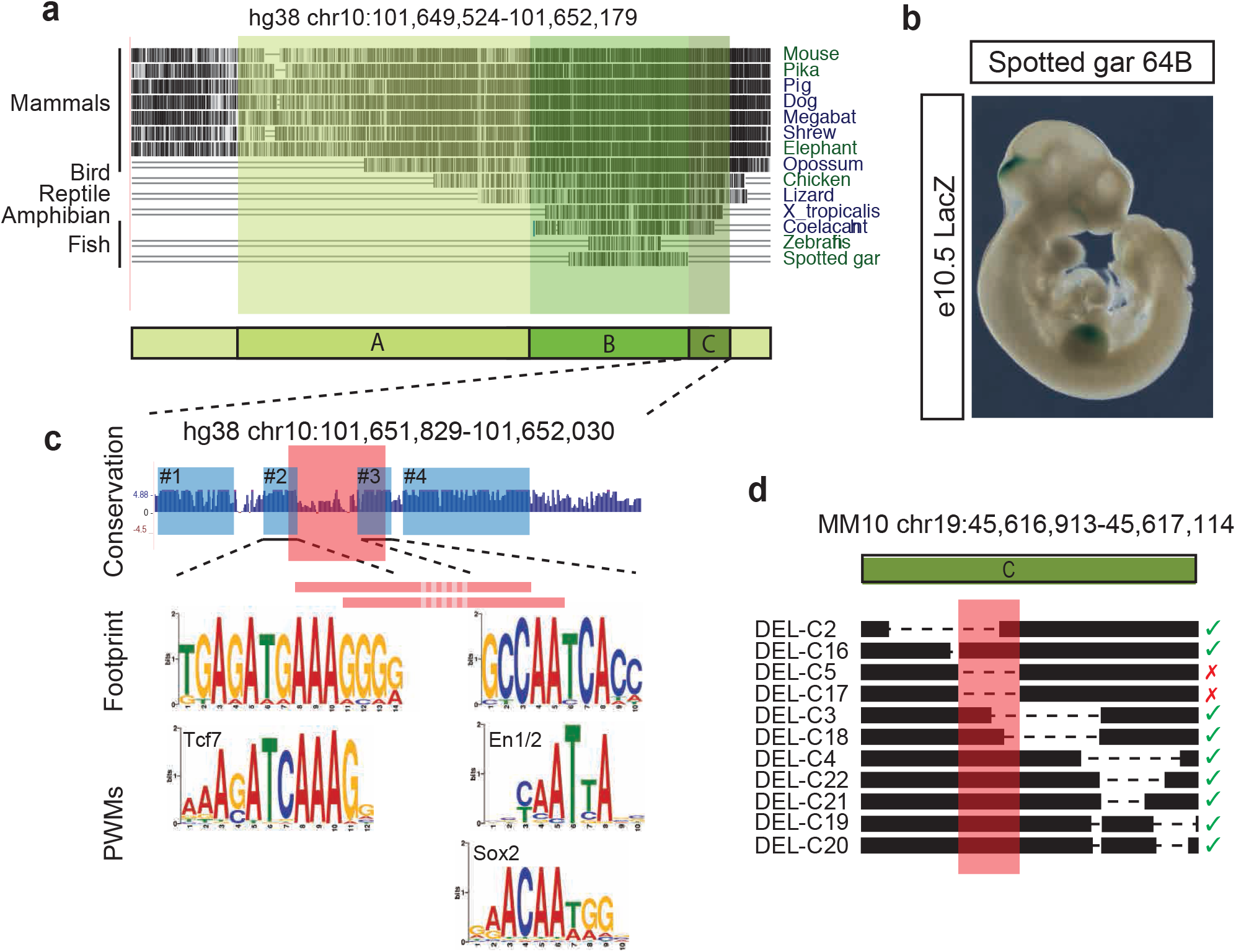
Cross-species comparison reveals non-conserved essential features of mouse of CE64. (**A**) Sequence conservation of CE64 across species. 64-B is conserved from fish to mammals while 64-C is conserved among tetrapods. (**B**) Photomicrograph of a transgenic embryo injected with a minimal reporter construct including spotted gar CE64 and stained for *LacZ* activity. Note that only 64-B is conserved in the spotted gar CE64. Arrowhead indicates the MHB. (**C**) Upper panel: sequence conservation score of 64-C. Blue boxes indicate highly conserved blocks. Red box indicate the smallest deletion that abrogates CE64 function. Middle panel: phylogenetic footprints generated from multiple sequence alignments corresponding to conserved block #2 and #3. The red bars indicate the breakpoints of the two smallest phenotype-causing deletions. Lower panel: PWMs of Tcf/Lef1, En1/2 and Sox proteins display similarities to the generated phylogenetic footprints. (**D**) Overview of small deletions in 64-C from the CRISPR/*Cas9* screen. Red box indicates 37bp depicted in (**C**). Red cross indicates loss of MHB derived tissues and green tick indicates normal brain morphology.

To investigate the sequence composition of CE64, we used multiple alignments to define phylogenetic footprints, hence identifying highly conserved sub-regions of the enhancer that might represent where functional TF binding can occur. In 64-B, the high conservation of the sequence precluded identification of obvious putative TFBS, while in 64-C we could define 4 conserved blocks of sequences resembling TFBS or TFBS clusters in length and composition (Fig6C, blue boxes, see alignments in FigS5). The 37bp deletion in 64-C abrogates two of these conserved blocks (red box in Fig6C and Fig6D), demonstrating that they are functionally important. Block #2 shares similarities with TCF/LEF binding sites (Fig6C), which can mediate responsiveness to Wnt-signalling, a known upstream inducer of *Fgf8* expression in the MHB ^34,35^. Noteworthy, mouse 64-B also comprises a potential Wnt-TCF/LEF response element (sequence CAGTTTCAAAGGAA). Block #3 bears homologies to the consensus binding motif defined for En1/2 (Fig6C), two transcription factors specifically expressed in the MHB ^36,37^ and that contribute to *Fgf8* maintenance there ^38^, as well to some extent to SOX proteins (Fig6C).

We then used these footprints to derive positional weight matrices (PWMs) and scan the spotted gar and zebrafish CE64 for corresponding motif occurrences. Only one of the two PWMs derived from the phylogenetic footprints in the 37bp deletion was detected in the spotted gar (block #3, Fig6C) or the zebrafish (block #2, Fig6C) CE64 (including the whole sequence tested in ref ^31^, and its spotted gar ortholog). These analyses suggest that although the orthologous CE64 elements that drive MHB expression in the teleost fishes and mammals may use different logics that could correspond to a rewiring of the *Fgf8* regulatory circuit.

## Discussion

Shadow and distributed enhancers have been described as common features in the regulatory genome that could provide robustness to gene expression, by buffering it against environmental changes and possible genetic variation ^39,40^. The *Fgf8* regulatory landscape is a prototypical example of the complexity of developmental gene regulation, which involves multiple enhancers with similar activity. By dissecting their function *in vivo* we found different acting logics within two sets of tissue-specific enhancers. In the limb, *Fgf8* AER expression results from the collective action of several enhancer modules with redundant activity (Fig7A). Noteworthy, the degree of conservation of the different AER enhancer does not seem to correlate with relative importance, and we do not see evidence that the “tetrapod” modules contribute specifically to the heterochronic shift associated with evolution of the AER from a primitive apical ectodermal fold ^41^. Contrarily to a simple view, the progressive recruitment of new AER enhancer modules during tetrapod evolution did not simply reinforce expression by addition of accessory elements to an ancestral essential enhancer. It may have allowed a redistribution of functional roles between the new elements, enabling more complex rewiring of the expression control of this gene in the apical ectoderm of the limb, which could have contributed to a prolonged maintenance of the apical ectodermal ridge, an essential step in the evolution of tetrapod limbs ^41,42^.

**Fig7.**
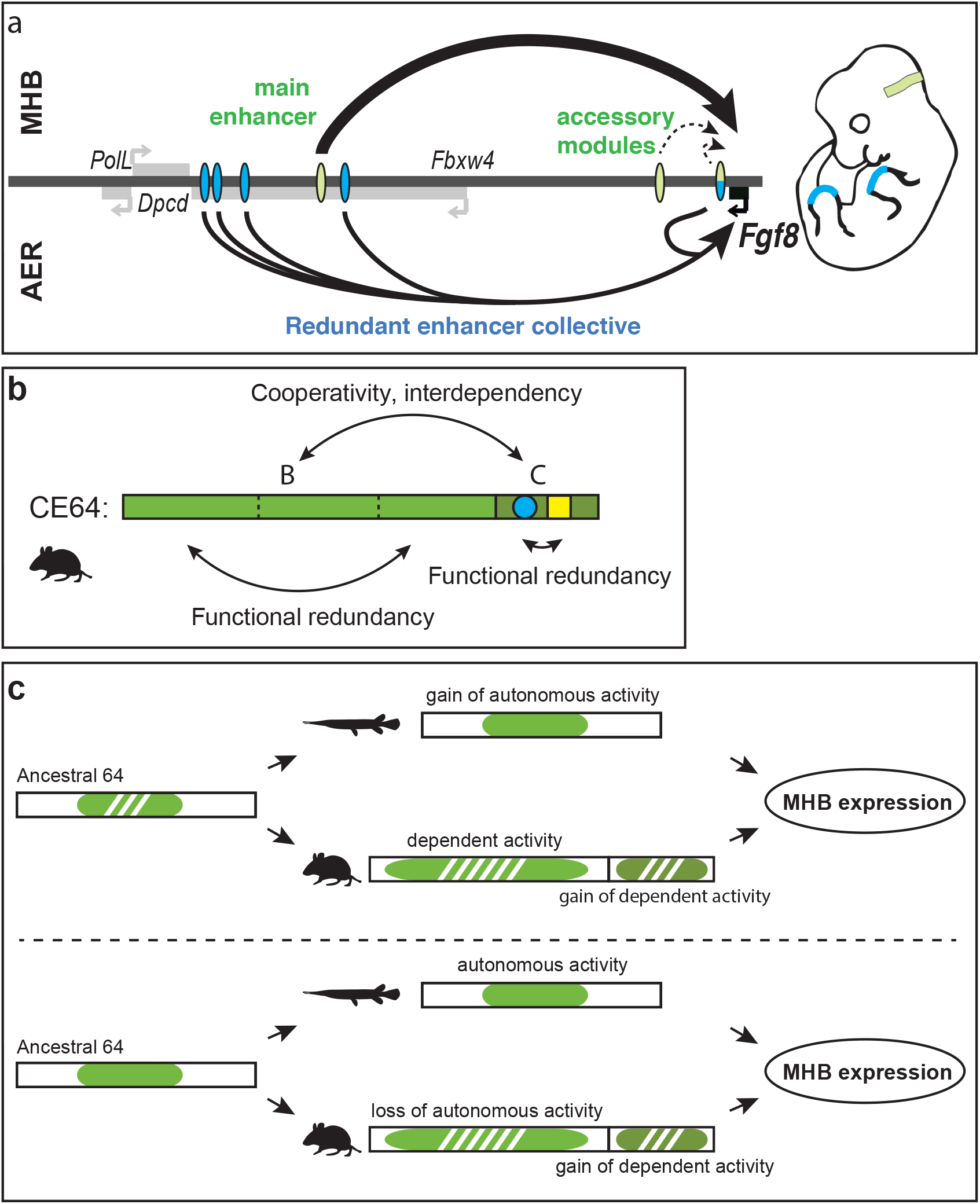
Two modes of regulatory redundancy provide robustness to *Fgf8* gene expression. (**A**) Schematic representation of enhancer activity in the MHB and the AER. In the MHB, one main enhancer is required and sufficient to direct *Fgf8* expression (upper panel). In the limb AER, a collective of redundant enhancers, each by themselves dispensable, directs gene expression (lower panel). Blue ovals: AER enhancers, green ovals: MHB enhancers (**B**) Schematic of the mouse CE64 and its subunits. CE64 is composed of two essential regulatory units that are reciprocally dependent and cannot alone direct expression in the MHB. Both of the regulatory subunits exert functional redundancy within themselves. This redundancy may be encoded by similar TFBS or recurrent DNA motifs in 64-B, while in 64-C it is encoded by two distinct DNA signatures (blue circle, yellow square) that reciprocally can buffer the loss of each other (**C**) Two scenarios for CE64 evolution. Upper panel: both spotted gar and the tetrapod lineage independently gained MHB regulatory activity, spotted gar within the ancestral enhancer (64-B in mouse) and tetrapods by addition of a new subunit (64-C). Lower panel: spotted gar CE64 retain an autonomous MHB regulatory activity in the absence of 64-C while in the tetrapod lineage, 64-B appears to have lost its autonomous activity and 64-C has been added to the regulatory wiring. Green area represents autonomous regulatory activity and dashed green non-autonomous activity.

In the MHB, *Fgf8* expression is solely dependent on one enhancer and the others appear dispensable (Fig7A). Given the high conservation of CE79 and CE80 and their previous identification also as putative enhancers in the MHB in the zebrafish ^29–31^, the finding that both are dispensable for normal development of the MHB region may be surprising. Still, it remains to be defined if those enhancers have important roles in other embryonic structures or later stages, and whether they may contribute to aspects of MHB function that cannot be assessed in a laboratory set-up (e. g. robustness to genetic or environmental variation).

Thorough dissection of the functional units of the essential MHB enhancer CE64 reveals a multi-layered organization, with separate units critical for its activity (Fig7B). The failure to initiate expression when deleting either of these regions demonstrates that their activities are interdependent (Fig7B). Our extensive *in vivo* screen of smaller deletions within CE64 nonetheless suggests that this enhancer can withstand relatively large sequence modifications, even in its evolutionary conserved parts. The small 37bp region, which deletion completely abrogates the function of the main MHB enhancer, thus causing loss of midbrain and cerebellum, identifies an essential and compact part of this enhancer. As removing overlapping bits of these 37bp does not lead to any phenotype, it demonstrates that redundancy is encoded in the regulatory architecture of the enhancer even at this scale and that most likely two sets of factors are involved (Fig7B). Sequence analysis suggests that Wnt-mediators LEF/TCF and En1/2 or Sox may be transcription factors associated with this activity.

Even though CE64 is highly conserved (to a degree that one could expect for an enhanceasome ^43^), the deletion analysis shows that it maintains function despite substantial sequence changes in key elements of its organisation, which fits better with a flexible “billboard” model of enhancer logic ^44,45^. Furthermore, the comparison of CE64 elements from different species argues indeed that its activity can use different logic in distinct animals. One of the two critical regions is only present in tetrapods, suggesting that it has recently evolved. This implies that the interdependence between 64-B and 64-C may have been acquired late during tetrapod evolution and may correspond to a change in *Fgf8* regulation. The fact that CE64 from spotted gar, in contrast to mouse 64-B, can drive expression in the MHB boundary of the mouse suggests that the spotted gar subunit gained regulatory potential or that some regulatory potential has been lost in the mouse enhancer subunit (Fig7C). We suggest that the addition of 64-C in the tetrapod lineage may have provided an additional layer of regulation that allowed for loss of ancestral features in 64-B, and eventually led to regulatory rewiring (Fig7C). Altogether, the dissection of 64 shows that it follows a complex logic involving multiple elements, which can both contribute to set up the very specific expression pattern of *Fgf8* in a given species, but as well allows for functional changes in its outcome on evolutionary timescales.

The complexity of developmental regulatory ensembles has made functional studies difficult. Here we demonstrate that Crispr/Cas9 *in vivo* deletion-screens can be very efficient in functionally dissecting their components and address this type of complexity. If several high-throughput screens have been conducted in cell lines using CRISPR/*Cas9* ^46–52^ our study is the first large one carried out *in vivo* in mouse embryos. The use of a large deletion to perform the screen in hemizygous conditions is not mandatory, but provide both increased yield and facilitate analysis. By focussing on function and not on activity, our approach provides an important complement to the transgenic enhancer bashing that has been performed so far, enabling to narrow down the essential sequences required for *Fgf8* expression *in vivo*. Such an approach is particularly necessary, given the intricate interplay between different units or enhancer modules, both at large scale within an ensemble and within an enhancer. Our study demonstrates the feasibility and usefulness of such approaches to decipher the complex, flexible and multi-scale organisation of developmental gene regulatory ensembles.

## Methods

### Animals and genotyping

All animal procedures were performed according to principles and guidelines at the EMBL Heidelberg (Germany) and the Institute Pasteur (Paris, France). Genotyping was performed by PCR using primers flanking deletion breakpoints (Supplementary methods TableS1). The breakpoints for all F1 pups of stable lines and all F0 embryos from the embryonic screen were sequenced. For the embryonic screen, primers internal to each deletion were used to identify any mosaic embryos carrying both deletion and wild type alleles. For some very small deletions, surveyor assays were used in addition to PCR to exclude mosaicism. The balancer mouse strains DEL(P-F8) and *Fgf8*^null^ were genotyped as previously described ^6^.

### Targeted genome engineering, in vitro fertilization and embryo transfers

Two CRISPR gRNA targets flanking each region of interest were designed using the CRISPR Design Tool (Zhang Lab, MIT) and are listed in Supplementary methods TableS2. *In vitro* transcription and cytoplasmic injections were performed essentially as described previously ^53^. *Cas9* from p×330 (Addgene) was subcloned downstream the T7 promoter in a pGEMte plasmid. The target plasmid was linearized, gel purified and used as template for IVT. Templates for gRNAs were generated through PCR amplification. IVT was performed with mMESSAGE mMACHINE T7 ULTRA kit (Life Technologies) and MEGAshortscript T7 kit (Life Technologies), respectively, and RNA was purified using MEGAclear kit (Life Technologies). *Cas9* mRNA (100ng/ul) and chimeric gRNAs (50ng/ul) were diluted in microinjection buffer ^54^ and injected according to standard procedure. For deletion screening of embryos, *in vitro* fertilisation (IVF) was performed the night before injections. One DEL(P-F8) heterozygous male was euthanized, the epididymis was dissected out and incubated 25-45 minutes in fertiup medium at 37°C, 5% CO_2_, allowing sperm to swim out. Meanwhile, oocytes from superovulated females were isolated into 200ul CARD media and 10-20 ul sperm was added before incubation over night.

### Cloning, transgenesis and X-gal staining

Transgenesis was performed as previously described ^6^. Briefly, fragments of interest were cloned upstream a ß-globin-derived minimal promoter and a *LacZ* reporter gene. The Tg(DEL-B) and Tg(DEL-C) fragments were cloned from CRISPR-embryo DEL-AB-2 and DEL-C respectively. Primers used for cloning are listed in TableS3. Linearized and gel-purified fragments were microinjected into fertilized mouse oocytes and transferred to pseudo-pregnant females (Institute Pasteur, Mouse Genetics Engineering). Embryos were collected at e10.5 and stained for ß-galactosidase activity using standard protocol. Genotyping PCR was performed on yolk sac DNA.

### Optical projection tomography

Embryonic brains were dissected free at e18.5, fixed in 4% PFA O/N and prepared for OPT scanning ^55^. Each specimen was scanned using the Bioptonics 3001 OPT scanner with a resolution of 1024×1024 pixels and reconstructed with the NRecon version 1.6.9.18 (Skyscan) software. Post-acquisition alignment values for reconstructions were calculated using LLS-Gradient based A-value tuning ^56^. Screenshots were exported from OPT volume renderings generated in Drishti v2.6.3 ^57^ and processed in Photoshop CS5 version 9.0.2 (Adobe). All image adjustments were applied equally to entire images and occasional artefacts such as fibers or dust were digitally removed.

### Gene expression analysis

In situ hybridisation was performed according to standard protocols with previously published *Fgf8* probe ^58^. For RT-qPCR, the MHB-region was dissected from e10.5 embryos and total RNA was extracted using the RNAeasy (Qiagene) kit. cDNA was prepared using the ProtoScript First Strand cDNA Synthesis Kit (New England Biolabs) with random primers. Fore each reaction, 150-200ng RNA was used. RT-PCR was performed according to manufacturers protocol on a GE48.48 IFC (Fluidigm) using SsoFast EvaGreen Supermix with low ROX (Fluidigm). Before RT-PCR, 10 (MHB) or 14 (limb) cycles of preamplification (Fluidigm PreAmp Master Mix) was performed using 15ng of input cDNA. Preamplified DNA was diluted 5 (MHB) or 10 (limb) times before RT-PCR reaction. Primers used are listed in (Supplementary methods TableS4).

### Motif analysis

For phylogenetic footprints, sequences of interest were retrieved from pre-calculated alignments at UCSC or Ensembl genome browsers; realigned using MUSCLE, and PWMs were calculated from these alignments. Motif analysis was performed using the online interface of the MEME suite ^59^.

## Acknowledgements

We are grateful to EMBL Genecore as well as to the Technology Core of the Center for Translational Science (CRT) at Institute Pasteur for support in conducting this study. We thank the DKFZ light microscopy facility and F. Bestvater for access to optical projection tomography equipment. We thank I. Braasch and H. Marlow for providing spotted gar tissue and extracted DNA, respectively. We thank greatly Y. Petersen (EMBL transgene facility) and all members of the animal facilities at both EMBL Heidelberg and Institute Pasteur for their help.

## Author contributions

F.S conceived the project and A.H. and F.S. designed the experimental strategies. A.H. performed or supervised all experiments. K.L. and S.B contributed to mouse embryos injections and transfers, and *in situ* hybridisation and skeletal preparations, respectively. F.L. produced DEL-B and DEL-C mutant mouse lines as well as transgenic LacZ reporter embryos; A.H. and F.S. wrote the paper with input of all authors.

