## Supplementary information for "Dissection of the *Fgf8* regulatory landscape by *in vivo* CRISPR-editing reveals extensive inter- and intra-enhancer redundancy"

Hornblad et al.

### Supplementary information

Supplementary Figures S1-S5

Supplementary Tables S1-S6

Supplementary Videos S1-S4

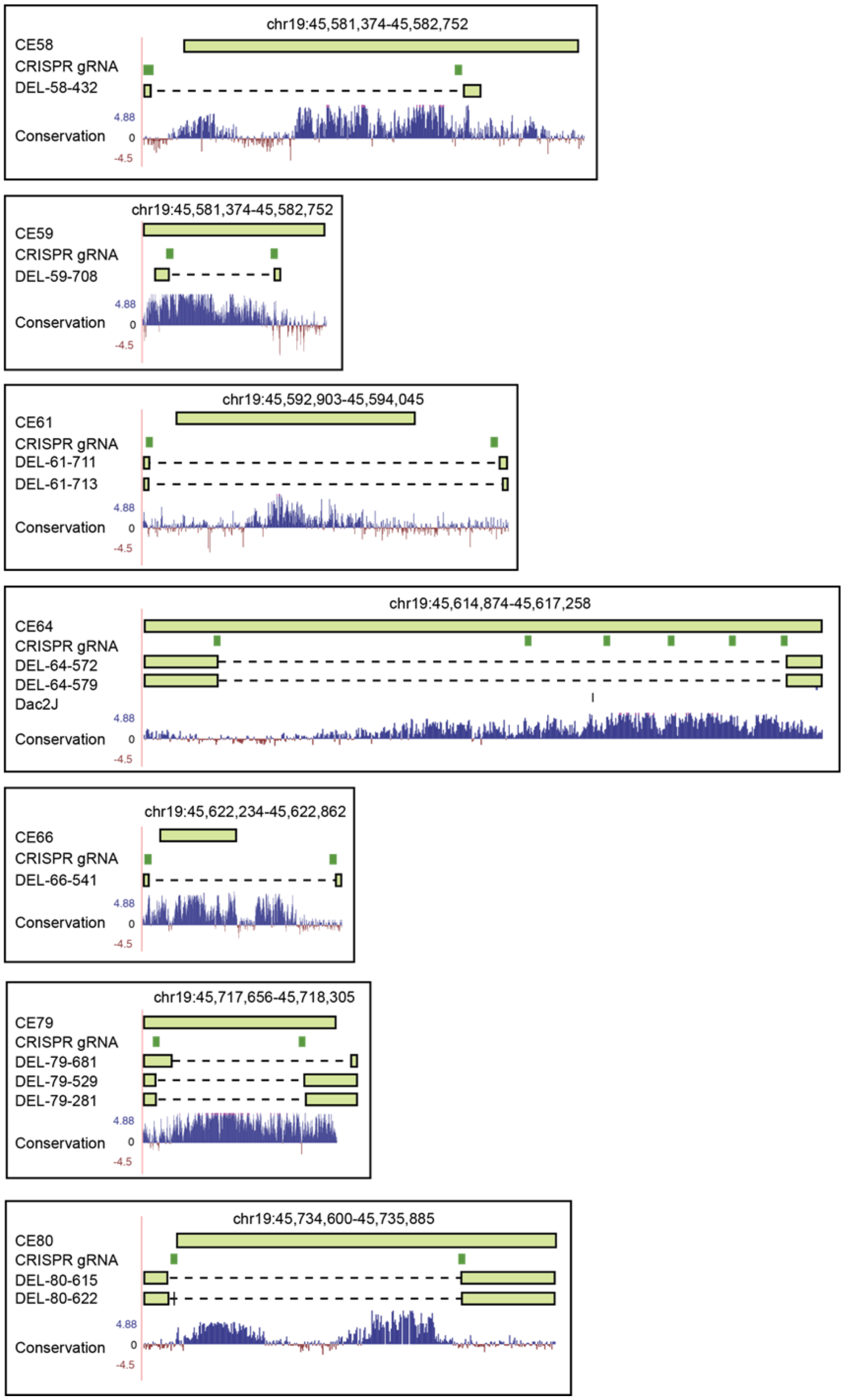


**FigS1. Genomic coordinates of the seven putative enhancers.** Upper row depicts mouse sequences homologous to human sequences with reported enhancer activity. Green boxes indicate CRISPR gRNAs used to create enhancer deletions. Dashed lines depict the location of the deletions for all lines used. DEL79-281 indicates the deletion of the DEL79-80 line that was generated over DEL80-622.

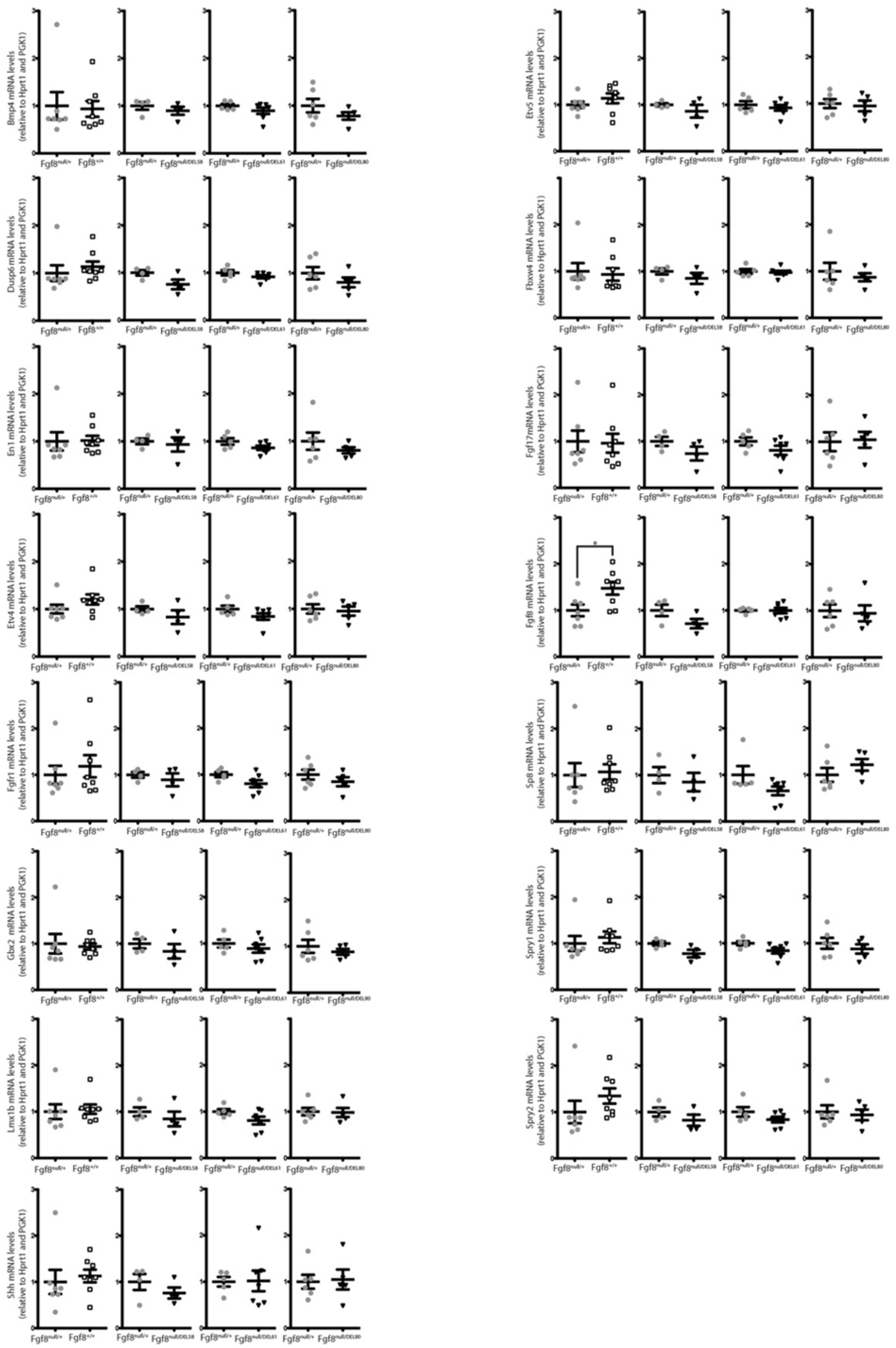

**FigS2. Gene expression levels are maintained in AER enhancer mutants.** RT-qPCR analysis of e10.5 forelimb in *Fgf8*$^{null/+}$, WT (n=7), DEL58 (n=4), DEL61 (n=5) and DEL80 (n=5) embryos. Relative expression of indicated mRNA as compared to heterozygous *Fgf8*$^{null/+}$ littermate controls. Individual data points as well as mean ± SEM are indicated. *p<0.05 (two-tailed Student's *t*-test).

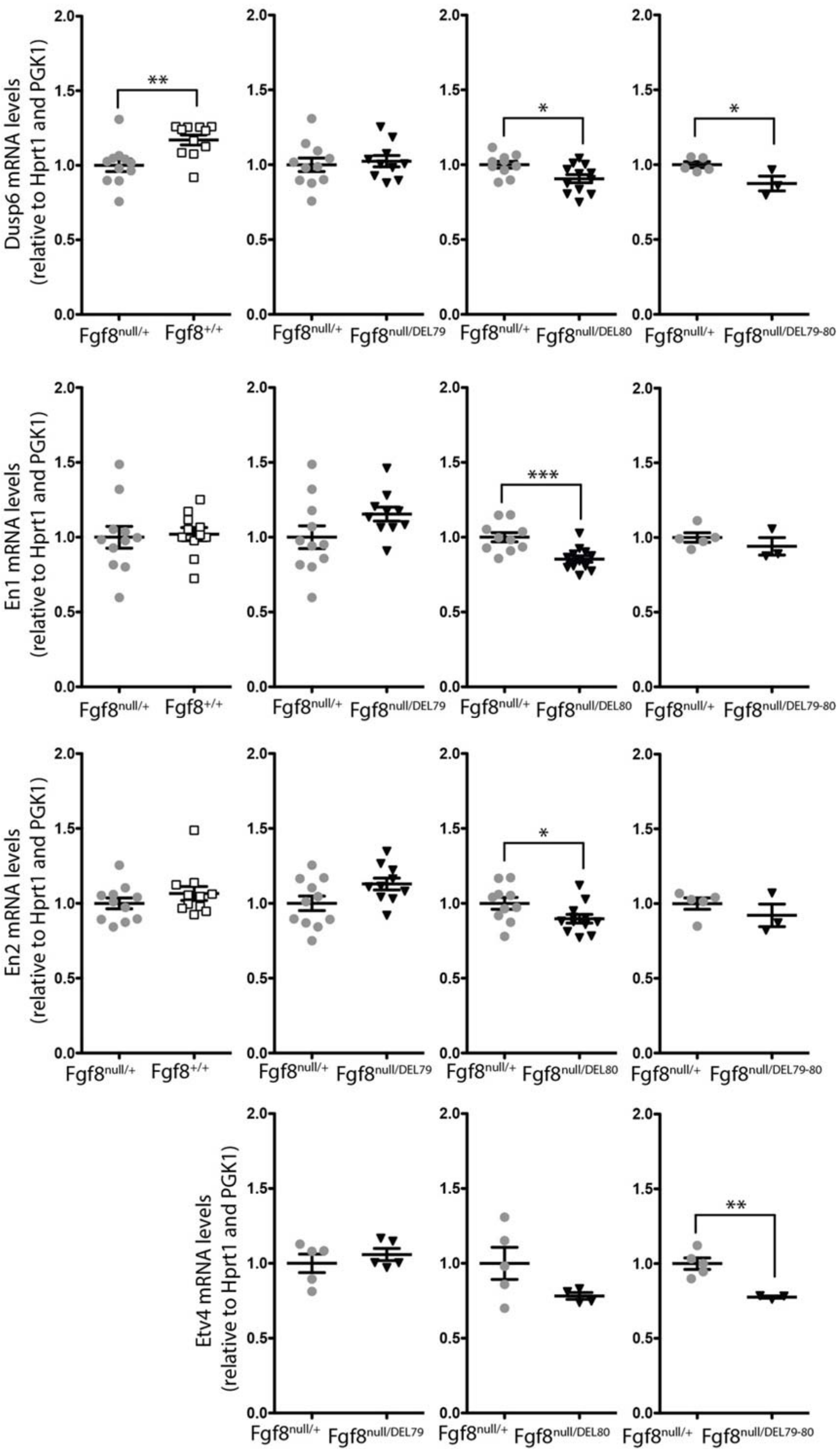


**FigS3. Minor changes in gene expression levels accompany DEL79. DEL80 and DEL79-80 mutants.** RT-qPCR analysis of e10.5 dissected MHB region in $Fgf8^{null/+}$ (n=11, 5/11, 5/10, 5), WT (n=11) , DEL79 (n=5/10), DEL80 (n=4/12) and DEL79-80 (n=3) embryos. Relative expression of indicated mRNA as compared to heterozygous $Fgf8^{null/+}$ littermate controls. Individual data points as well as mean ± SEM are indicated. *p<0.05, **p<0.01, ***p<0.001 (two-tailed Student's *t*-test).

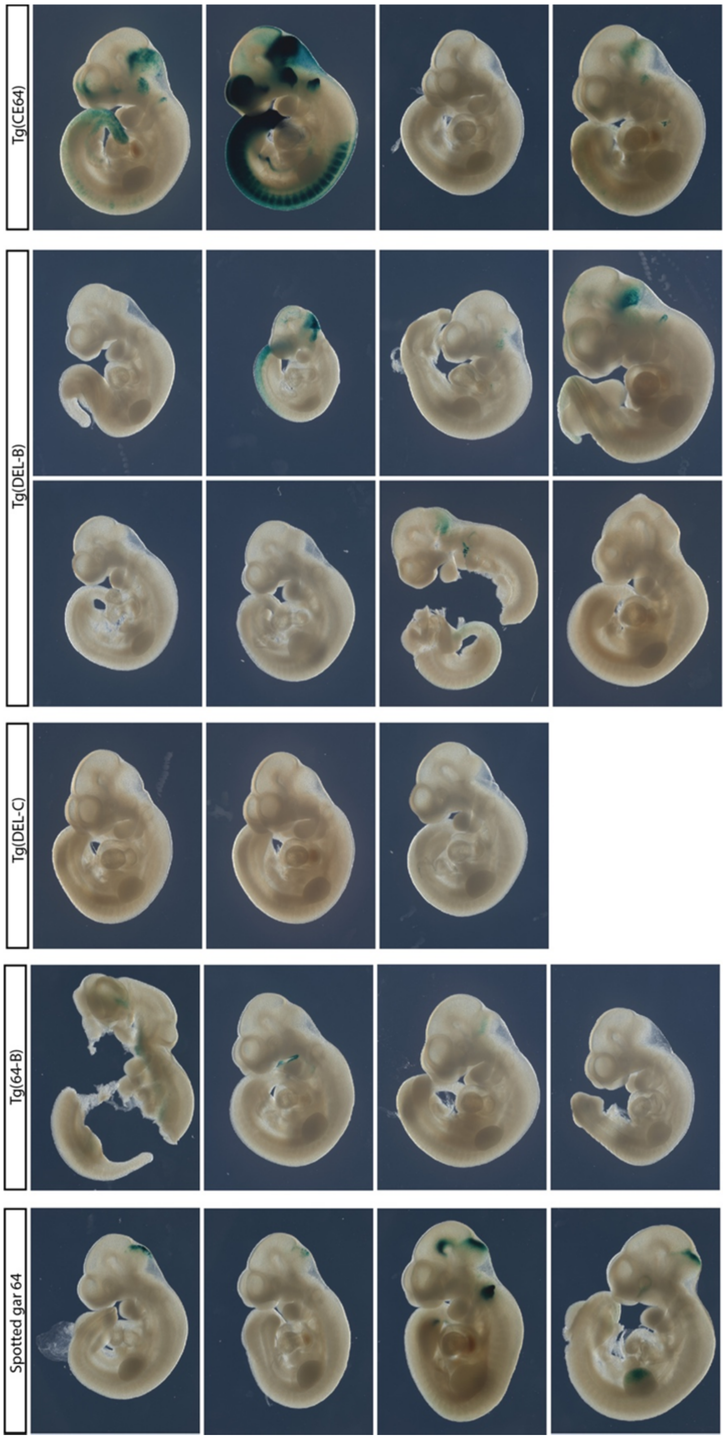


**FigS4. Transgenic enhancer activity of mouse CE64 and spotted gar CE64.** Compilation of all transgenic embryos harvested for Tg(CE64), Tg(DEL-B), Tg(DEL-C), Tg(64-B) and Tg(Spotted gar64B) . Expression is absent in the MHB of Tg(DEL-B), Tg(DEL-C) and Tg(64-B) embryos while all Tg(Spotted gar 64B) embryos display expression in the MHB region. Note also the hindbrain expression present in 4 of the Tg(DEL-B) embryos.

```
Block#1
Human                      GGACAACAAGGAAAATCGGCTG-GCCCATTGT
Mouse                      GGACAACAAGGAAAATCGGCTG-GTCCATTGT
Pika                       GGACAACAAGGAAAATCGGCTG-GCCTATTGT
Pig                        GGACAACAAGGAAAATCGGCTG-GCCCATTGT
Dog                        GGACAACAAGGAAAATCGGCGG-GCCCATTGT
Megabat                    GGACAACAAGAAAAATCGGCTG-GCCCATTGT
Shrew                      GGACAACAAGGAAAATCGGCCA-GCCCATTGT
Elephant                   GGACAACAAGGAAAATCGGCTG-GCCCATTGT
Opossum                    GGACAACAAGGAAAATAGGCTG-GCCCATTGT
Chicken                    GAACAACAAGGAAAATATGCTG-GCCCATTGT
Lizard                     GAACAACAAGGAAAATATGCTGAGTCCATTGT
Xenopus                    AAACAACAAGAAAAATCTGCT--TTCTATTGT
Coelacanth                 AAGCAACAAGAAAAACCTGCT--ACCCATTGT
Zebrafish                  ----------------------GCTCATTGC
Spotted_gar   -------------------------------

Block#2
Human                      GTGCTAAGTGAGATGAAAGGGG
Mouse                      GTGCTAAGTGAGATGAAAGGGG
Pika                       GTGCTAAGTGAGATGAAAGGGG
Pig                        GTGCTAAGTGAGATGAAAGGGG
Dog                        GTGCTAAGTGAGATGAAAGGGA
Megabat                    GGTGCTAAGTGAGATGAAAGGG
Shrew                      GTGCTAAGTGAGATGAAAGGGG
Elephant                   GTGCTAAGTGAGATGAAAGGGG
Opossum                    GTGCTAAGTGAGATGAAAGCAG
Chicken                    GCACCAAGGGAGATGAAAGGGA
Lizard                     ACACTATAGGAGATGAAAGGGA
Xenopus                    GTACTAAGTGAGATGAAAGGGA
Coelacanth                 GAGCAGGATTAAAAAAAAAGGG
Zebrafish                  --GCT-----------------
Spotted_gar   ----------------------

Block#3
Human                      CCCGCCTGCCAATCA
Mouse                      CCCGCCTGCCAATCA
Pika                       CCCGCCTGCCAATCA
Pig                        CCCGCCTGCCAATCA
Dog                        CCCGCCTGCCAATCA
Megabat                    CCCGCCTGCCAATCA
Shrew                      CCCGCCTGCCAATCA
Elephant                   CCCGCCTGCCAATCA
Opossum                    CCCGCCTGCCAATCA
Chicken                    ACCACCTGCCAATCA
Lizard                     GCTGCCTGCCAATCA
Xenopus                    ACAGCCTGCCAATCA
Coelacanth                 CCATCCTGTCAATCA
Zebrafish                  ---------------
Spotted_gar                -CCACCTCCCAACCA

Block#4
Human                      AGTTATT-AAGGTCACAATGTTAAATTCTTTTGACCTGCACCTAATCAAAGAT
Mouse                      AGTTATT-AAGGTCACAATGTTAAATTCTTTTGACCTGCACCTAATCAAAGAT
Pika                       AGTTATT-AAGGTCACAATGTTAAATTCTTTTGACCTGCACCTAATCAAAGAT
Pig                        AGTTATT-AAGGTCACAATGTTAAATTCTTTTGACCTGCACCTAATCAAAGAT
Dog                        AGTTATT-AAGGTCACAATGTTAAATTCTTTTGACCTGCACCTAATCAAAGAT
Megabat                    AGTTATT-AAGGTCACAATGTTAAATTCTTTTGACCTGCACCTAATCAAAGAT
Shrew                      AGTTATT-AAGGTCACCATGTTAAATTCTTTTGACCTGCACCTAATCAAAGAT
Elephant                   AGTTATT-AAGGTCACAATGTTAAATTCTTTTGACCTGCACCTAATCAAAGAT
Opossum                    AGTTATT-AAGGTCACAATGTTAAATTCTTTTGACCTGCACCTAATCAAAGAT
Chicken                    AGTTATT-AAGGTCACGATGTGAAATTCTTTTGACCTACACCTAATCAAAGAT
Lizard                     ACTTATT-AAGGTCACAATGT-AGACAATATTAACCTGAATCCAATCAAAGAT
Xenopus                    AGTAAGCAAAGGTCACAATGTTACATTCTTTTGACCTGCACTTA---------
Coelacanth                 AGGTATC-AA-------------------------------------------
Zebrafish                  -----------------------------------------------------
Spotted_gar   -----------------------------------------------------
```

**FigS5. MUSCLE alignments of conserved blocks in 64-C.** Disruption of both block #2 and #3 together through CRISPR/Cas9 editing cause absence of MHB-derived structures while deletion of any single block does not cause phenotype.

**TableS1.** Genomic coordinates of the CRISPR deletions for all founders used in this study.

| Strain name | Target CE | Genomic coordinates of deletions MM10 | Size of deletion (bp) |
| --- | --- | --- | --- |

| | | | |
|---|---|---|---|
| 58-432 | CE58 | 45,581,399 - 45,582,371 | 973 |
| 59-708 | CE59 | 45,585,002 - 45,585,325 | 324 |
| 61-711 | CE61 | 45,592,927 - 45,594,008 | 1082 |
| 61-713 | CE61 | 45,592,924 - 45,594,025 | 1102 |
| 64-572 | CE64 | 45,615,282 - 45,617,095 | 1811 |
| 64-579 | CE64 | 45,615,283 - 45,617,093 | 1814 |
| 66-541 | CE66 | 45,622,255 - 45,622,840 | 586 |
| 79-529 | CE79 | 45,717,702 - 45,718,193 | 492 |
| 79-681 | CE79 | 45,717,759 - 45,718,349 | 591 |
| 79-281 | CE79 | 45,717,706 - 45,718,198 | 493 |
| 80-622 | CE80 | 45,734,600 - 45,735,569 | 970 |
| 80-615 | CE80 | 45,734,594 - 45,735,569 | 976 |

**TableS2.** Genomic coordinates, type of modification and phenotypic outcome for all embryonic CRISPR deletions reported in this study.

| Name | Coordinates MM10 | Size | Intact MHB | Type of modification |
|---|---|---|---|---|
| DEL-AC1 | 45,615,244 - 45,617,008 | 1765 | ✗ | Deletion |
| DEL-A1 | 45,615,265 - 45,616,287 | 1023 | ✓ | Deletion |
| DEL-AB1 | 45,615,282 - 45,616,929 | 1648 | ✗ | Deletion |
| DEL-AB3 | 45,615,282 - 45,616,932 | 1651 | ✗ | Deletion |
| DEL64 | 45,615,283 - 45,617,093 | 1814 | ✗ | Deletion |
| DEL-AB2 | 45,615,790 - 45,616,928 | 1139 | ✗ | Deletion |
| DEL-B1 | 45,616,275 - 45,616,928 | 654 | ✗ | Deletion |
| DEL-B8 | 45,616,221 - 45,616,929<br>45,617,090- 45,617,107 | 709<br>18 | ✗ | Deletion |
| DEL-B2 | 45,616,273 - 45,616,516 | 244 | ✓ | Deletion |
| DEL-B4 | 45,616,517 - 45,616,722 | 206 | ✓ | Deletion |
| DEL-B3 | 45,616,517 - 45,616,928 | 412 | ✓ | Deletion |
| DEL-BC1 | 45,616,542 - 45,617,093 | 552 | ✗ | Deletion |
| DEL-BC2 | 45,616,672 - 45,617,098<br>45,616,100 - 45,617,257 | 427<br>159 | ✗ | Deletion<br>Inversion |
| DEL-B7 | 45,616,720 - 45,616,930 | 211 | ✓ | Deletion |
| DEL-BC4 | 45,616,721 - 45,618,022 | 1302 | ✗ | Deletion |
| DEL-B6 | 45,616,722 - 45,616,936 | 215 | ✓ | Deletion |
| DEL-B5 | 45,616,726 - 45,616,929 | 204 | ✓ | Deletion |
| DEL-BC5 | 45,616,768 - 45,617,102 | 335 | ✗ | Deletion |
| DEL-C1 | 45,616,922 - 45,617,092 | 171 | ✗ | Deletion |
| DEL-C6 | 45,616,923 - 45,616,929 | 7 | ✓ | Deletion |
| DEL-C7 | 45,616,926 - 45,617,534 | 609 | ✗ | Deletion |
| DEL-C8 | 45,616,926 - 45,617,534 | 609 | ✗ | Deletion |
| DEL-C9 | 45,616,927 - 45,617,023 | 97 | ✗ | Deletion |
| DEL-C11 | 45,616,930 - 45,616,938<br>45,616,956 - 45,617,056 | 304 insertion + 9 deletion<br>101 | ✗ | Indel<br>Deletion |
| DEL-C2 | 45,616,930 - 45,616,995 | 66 | ✓ | Deletion |
| DEL-C10 | 45,616,930 - 45,617,055 | 126 | ✗ | Deletion |
| DEL-C12 | 45,616,930 - 45,617,091 | 162 | ✗ | Deletion |
| DEL-C13 | 45,616,930 - 45,617,094 | 165 | ✗ | Deletion |
| DEL-C15 | 45,616,935 - 45,617,257 | 323 | ✗ | Deletion |
| DEL-C16 | 45,616,967 - 45,616,971 | 5 | ✓ | Deletion |
| DEL-C17 | 45,616,968 - 45,617,005 | 38 | ✗ | Deletion |
| DEL-C5 | 45,616,971 - 45,617,007 | 37 | ✗ | Deletion |
| DEL-C3 | 45,616,991 - 45,617,056 | 66 | ✓ | Deletion |
| DEL-C18 | 45,616,999 - 45,617,055 | 57 | ✓ | Deletion |
| DEL-C4 | 45,617,045 - 45,617,146 | 102 | ✓ | Deletion |
| DEL-C20 | 45,617,051 - 45,617,056<br>45,617,088 - 45,617,112 | 6<br>25 | ✓ | Deletion<br>Deletion |
| DEL-C19 | 45,617,051 - 45,617,056<br>45,617,090 - 45,617,108 | 6<br>19 | ✓ | Deletion<br>Deletion |
| DEL-C21 | 45,617,056 - 45,617,081 | 26 | ✓ | Deletion |
| DEL-C22 | 45,617,056 - 45,617,094 | 39 | ✓ | Deletion |

**TableS3.** Primers used for genotyping.

| Primer name | Sequence |
|---|---|
| CE58R-CRISP1F-Surveyor | ACAACCAGGCTGTCTTCCAG |
| CE58R-CRISP1R-Surveyor | CTCCACCCTACCCCAAGTCT |
| CE58R-CRISP2F-Surveyor | TGTAGTGCTGTGGTGCTGAG |
| CE58R-CRISP2R-Surveyor | AGACTCAGGAGGCTAGGGTG |
| CE59-CRISP1F-Surveyor | AGAGCACACGTGTTCAGACA |
| CE59-CRISP1R-Surveyor | GGACTGCCCCTCTTGAAAGT |
| CE59R-CRISP2F-Surveyor | TCAGTTTGGTGTCAGCAGGC |
| CE59R-CRISP2R-Surveyor | GAACGTGGCTTCAGCTTGTG |
| CE61-CRISP1F-Surveyor | CTGTGGGAAGGATCGGTCTG |
| CE61-CRISP1R-Surveyor | TTTGGAGATGACAGGTGGGC |
| CE61-CRISP2F-Surveyor | ACCATCTGCCACCGAGAATG |
| CE61-CRISP2R-Surveyor | GAAGAAGGCAGGCACAAAGC |
| CE64-CRISP1F-Surveyor | AGAGTGACTGGCATCAGTGC |
| CE64-CRISP1R-Surveyor | GTAGAGGGAAGCATTGGGGG |
| CE64-CRISP2F-Surveyor | CCTGGGAAAAATGCAGGCAC |
| CE64-CRISP2R-Surveyor | GAGACCCAGTCCTGACCTCT |
| CE64-CRISP3F-Surveyor | TTTGGACACACTGACAGGGG |
| CE64-CRISP3R-Surveyor | GAGGAGGGCGGAATGAAGAG |
| CE66-CRISP1F-Surveyor | CCAAGGGAACCAAGTGTGGA |
| CE66-CRISP1R-Surveyor | AGGGGAAAGGGGAGCCATAA |
| CE66-CRISP2F-Surveyor | TTATGGCTCCCCTTTCCCCT |
| CE66-CRISP_R2-Surveyor | AGAGGACACAAACAGACGGG |
| CE79-CRISP1F-Surveyor | CACCAGTCCATGCAGACCAT |
| CE79-CRISP1R-Surveyor | TCACACACCATACCCCCTGA |
| CE79-CRISP2F-Surveyor | AATCTCCCTGAGAGTGGCCT |
| CE79-CRISP2R-Surveyor | ACCTCATTTCCCTGAGGGGT |
| CE80-CRISP1F-Surveyor | TAAGCATATTGGGCCGGCAA |
| CE80-CRISP1R-Surveyor | AGGTGAGCAGAAGAGAGGGT |
| CE80-CRISP2F-Surveyor | CCCAGCTCCTCGCACAATAA |
| CE80-CRISP2R-Surveyor | TTTTGGCACCTCTCTTGGCA |
| CE80-CRISP_F2-Surveyor | CCAAGACCTGTGTTGGGTCT |
| CE80-CRISP_R2-Surveyor | GAAGAGCAATTGCCCAGTGT |
| CE64-F1 | GACCTACACAGGCCGCATTA |
| CE64-R1 | AAGAGACAATGGACCAGCCG |
| CE64-F2 | TCTAGCAATTTGGAGGCGGG |
| CE64-R2 | AAGATTTGGCATGGGAGCCA |
| 64-F3 | CCTCGGCACGCCATCTATAC |
| CE64-R3 | AGATACATCGCAAGGCAGCA |
| Tcf/Lef1 DEL | TGCTAAGTGAGATGAAAGGGGCC |

**TableS4.** CRISPR gRNAs used for zygote injections.

| gRNA Name | CE Target | Sequence |
|---|---|---|
| CE58-CRISP1 | 58 | CCAAGCACCTTGGCCGTGGGAGG |
| CE58-CRISP2 | 58 | GGGCGGCGTGCGCAGAGCCTTGG |
| CE59-CRISP1 | 59 | TAACAAGGCGACATAATTACAGG |
| CE59-CRISP2 | 59 | CTCTCCCCCGCCCCGTACAGGGG |
| CE61-CRISP1 | 61 | GTACCCACCACGTCAGCGGGAGG |
| CE61-CRISP2 | 61 | CGGCAACAAGCCCGATCCATGGG |
| CE64-CRISP1 | 64 | TCAGAAGCTTTACCGCCTAGAGG |
| CE64-CRISP2 | 64 | TGCCACCTGGTACTTCCCGCAGG |
| CE64-CRISP3 | 64 | CCTTTCAAATCCGAATGGGGAGG |
| CE64-CRISP4 | 64 | GTATGATTTCACGTCAGAACAGG |
| CE64-CRISP5 | 64 | GAGGGAGAAGCAACCCCGATGGG |
| CE64-CRISP6 | 64 | GGACAACAAGGAAAATCGGCTGG |
| CE64-CRISP7 | 64 | CGGTAAAGCTTCTGAGAATCTGG |
| CE64-CRISP8 | 64 | CTCAGCTCTAACTGGGCCTCAGG |
| CE64-CRISP9 | 64 | GTGCATCTTTGATTAGGTGCAGG |
| CE64-CRISP10 | 64 | GTGATTGGCAGGCGGGCTATGGG |
| CE64-CRISP11 | 64 | GTGCATCTTTGATTAGGTGCAGG |
| CE64-CRISP12 | 64 | TGTGCTAAGTGAGATGAAAGGGG |
| CE64-CRISP13 | 64 | TTAGCACAGAGAAGAGACAATGG |
| CE64-CRISP14 | 64 | ACTAGAGGTGATTGGCAGGCGGG |
| CE64-CRISP15 | 64 | AATAACTAGAGGTGATTGGCAGG |
| CE64-CRISP16 | 64 | CCAATCACCTCTAGTTATTAAGG |
| CE66-CRISP3 | 66 | AACAACTCCCTTTGAACCTCCGG |
| CE66-CRISP6 | 66 | TACAGGGGCAGTGTAGTAATCGG |
| CE79-CRISP1 | 79 | TGCAGAGCAGCCCGGTAGCTGGG |
| CE79-CRISP2 | 79 | TATCACGGCCCGGAGCTCAGAGG |
| CE80-CRISP2 | 80 | CTTCAGGTGCGGGGACACGGGGG |
| CE80-CRISP6 | 80 | AAGGCAGACCACTCTATCCCTGG |

**TableS5.** Cloning primers used for transgenic constructs.

| Primer name | Sequence | Target construct |
|---|---|---|
| 64-1/3 F | TGaagcttCTAGAGGCAGAGCAGCTCAG | Tg(CE64), Tg(DEL-C), Tg(DEL-B) |
| 64-1/3 R | TGctcgagCCCCATTCGGATTTGAAAGGGCCGG | Tg(CE64), Tg(DEL-B) |
| 64-6/3DEL R | TGctcgagCCCCAGATTTTCCTTGTTGTCC | Tg(DEL-C) |
| 64-2/5 F | TGaagcttTCCCTCAGCGATTGCACAG | Tg(64-B) |
| 64-2/5 R | TGctcgagGATTTTCCTTGTTGTCCCTTGTTTG | Tg(64-B) |
| 64-LepOcu F | TGaagcttATGGATGTGGTTGGGAGGTG | Spotted gar 64 |
| 64-LepOcu R | TGctcgagGTCTCGCTGTACGCTTATACAAAG | Spotted gar 64 |

**TableS6.** RT-qPCR primers used for gene expression analysis and their corresponding targets.

| Primer name | Sequence | Target gene |
|---|---|---|
| #2579_Mm_RT_Dusp6_MKP3_Fwd | GAATGAGAACACTGGTGGAGAGTCG | Dusp6 |
| #2580_Mm_RT_Dusp6_MKP3_Rev | ACTCGGCCTGGAACTTACTGAAGC | |
| #2569_Mm_RT_En1_Fwd | TGGGTCTACTGCACACGCTATTCG | En1 |
| #2570_Mm_RT_En1_Rev | GCCGCTTGTCTTCCTTCTCGTTC | |
| #2555_Mm_RT_En2_Fwd | GCTATTCTGACCGGCCTTCTTCAG | En2 |
| #2556_Mm_RT_En2_Rev | GGTACCTGTTGGTCTGAAACTCAGC | |
| #2567_Mm_RT_Etv4_PEA3_Fwd | AGGAAGCCACCACTCCCCTACCAC | Etv4 |
| #2568_Mm_RT_Etv4_PEA3_Rev | GGGACTTGATGGCGATTTGTCTG | |
| RTqPCR_Etv5_Fwd | TGAGCAGTTTGTCCCAGATTTTCAG | Etv5 |
| RTqPCR_Etv5_Rev | CTGTGCAGCTCCCGTTTGATCTTG | |
| #2529_Mm_RT_Fbxw4 _Fwd | CCCACAACAAGCTGTTCCAGTCAC | Fbxw4 |
| #2530_Mm_RT_Fbxw4 _Rev | TACACACCATCCAGACCGAAGACC | |
| #2599_Mm_RT_FGF17_Fwd | AAGAGGGGCAAGCTGATTGGGAAG | Fgf17 |
| #2600_Mm_RT_FGF17_Rev | AAAGCCATGAACCAGCCCTCGTG | |
| #2601_Mm_RT_FGF18_Fwd | GGCGAGGACGGGGACAAGTATG | Fgf18 |
| #2602_Mm_RT_FGF18_Rev | CCGGACTTGACTCCCGAAGGTATC | |
| #2531_Mm_RT_Fgf8_Fwd | TATCGGTCTCCACAATGAGCTTCG | Fgf8 |
| #2532_Mm_RT_Fgf8_Rev | CCTGGCCAACAAGCGCATCAAC | |
| Fwd_SYBR_FgFR1 | TCTGGCCTCTACGCTTGC | Fgfr1 |
| Rev_SYBR_FgFR1 | AGGATGGGAGTGCATCTGA | |
| #2561_Mm_RT_Gbx2_Fwd | GCTCGCTGCTCGCTTTCTCTGC | Gbx2 |
| #2562_Mm_RT_Gbx2_Rev | GCTGTAATCCACATCGCTCTCCAG | |
| #2568_Mm_RT_Etv4_PEA3_Rev | GGGACTTGATGGCGATTTGTCTG | Bmp4 |
| #2569_Mm_RT_En1_Fwd | TGGGTCTACTGCACACGCTATTCG | |
| Hprt_mRNA_F | CTTCCTCCTCAGACCGCTTTT | Hprt |
| Hprt_mRNA_R | CATCATCGCTAATCACGACGC | |
| #2571_Mm_RT_Lmx1b_Fwd | AGATGAAGAAGCTGGCCCGGAGAC | Lmx1b |
| #2572_Mm_RT_Lmx1b_Rev | CTCCATGCGGCTTGACAGAACCTC | |
| #2557_Mm_RT_Otx2_Fwd | CTGCATCCCTCCGTGGCTACC | Otx2 |
| #2558_Mm_RT_Otx2_Rev | CGTCGAGCTGTGCCCTAGTAAATG | |
| #2507_Mm_RT_Pax2_Fwd | CGAGGGCATCTGCGATAATGAC | Pax2 |
| #2508_Mm_RT_Pax2_Rev | CTGCTGAACTTTGGTCCGGATG | |
| Fwd_SYBR_Pax5 | ACGCTGACAGGGATGGTG | Pax5 |
| Rev_SYBR_Pax5 | GGGGAACCTCCAAGAATCAT | |
| mPax6-qpcr-fwd | CAAACACACATGAACAGTCAGC | Pax6 |
| mPax6-qpcr-rev | ACTTGGACGGGAACTGACAC | |
| Fwd_SYBR_MmPGK1 | TACCTGCTGGCTGGATGG | PGK1 |
| Rev_SYBR_MmPGK1 | CACAGCCTCGGCATATTTCT | |
| #2565_Mm_RT_Sp8_Fwd | TCGACGATTTCGAAGGGGAACG | Sp8 |
| #2566_Mm_RT_Sp8_Rev | TGGGGCTGCCGATCTTATTACAGG | |

| | | |
|---|---|---|
| #2551_Mm_RT_Spry1_Fwd | GGTCATAGGTCAGATCGGGTCATCC | Spry1 |
| #2552_Mm_RT_Spry1_Rev | TGGGTGGGGTCCTCTTTCAAGG | |
| #2549_Mm_RT_Spry2_Fwd | AAGGGAGAGGGGTTGGTGCAAAG | Spry2 |
| #2550_Mm_RT_Spry2_Rev | CCATCAGGTCTTGGCAGTGTGTTC | |
| #2577_Mm_RT_Shh_Fwd | CCCCAATTACAACCCCGACATC | Shh |
| #2578_Mm_RT_Shh_Rev | GGCATTTAACTTGTCTTTGCACCTCTG | |
| #2559_Mm_RT_Wnt1_Fwd | CCAACAGTAGTGGCCGATGGTG | Wnt1 |
| #2560_Mm_RT_Wnt1_Rev | CAGACTCTTGGAATCCGTCAACAGG | |

**VideoS1. DEL64 mutants display severe hypoplasia of the midbrain and cerebellum.** Tomographic sections from e18.5 mutant brain generated by optical projection tomography. Note the absence of all the MHB-derived structures: the superior colliculus, inferior colliculus, isthmus, and cerebellum.

**VideoS2. DEL79 mutants display normal brain anatomy.** Tomographic sections from e18.5 mutant brain generated by optical projection tomography. Note the normal appearance of all MHB derived structures: the superior colliculus, inferior colliculus, isthmus, and cerebellum.

**VideoS3 DEL80 mutants display normal brain anatomy.** Tomographic sections from e18.5 mutant brain generated by optical projection tomography. Note the normal appearance of all MHB derived structures: the superior colliculus, inferior colliculus, isthmus, and cerebellum.

**VideoS4. DEL79-80 mutants display normal brain anatomy.** Tomographic sections from e18.5 mutant brain generated by optical projection tomography. Note the normal appearance of all MHB derived structures: the superior colliculus, inferior colliculus, isthmus, and cerebellum.
